# Disrupted FAP-satellite cell communication contribute to maladaptive muscle remodeling in older women

**DOI:** 10.64898/2026.09.23.753766

**Authors:** Chad M. Skiles, Zachary J. Fennel, Paul-Emile Bourrant, Elena M. Yee, Robert J. Castro, Alexander R. Keeble, Ryan M. O’Connell, Christopher S. Fry, Micah J. Drummond

## Abstract

Aged skeletal muscle has decreased ability to rebound from physiological stressors such as disuse atrophy. We previously observed that older adults during recovery following muscle disuse were characterized by rapid skeletal muscle immune cell expansion, cellular senescence and collagen deposition, responses that were biased toward older women. Here we used single-nucleus RNA sequencing and complementary *in vitro* muscle primary cell experiments in young (YF; 22±3y; n=8) and older females (OF; 66±5y; n=9) to investigate age-mediated cellular function and intercellular communication during recovery from disuse atrophy. Compared with YF, OF exhibited a markedly greater transcriptional response at 7d-recovery (YF: 1,548 DEGs, OF: 7,999 DEGs), driven primarily by slow- (YF: 96 DEGs, OF: 984 DEGs) and fast-twitch myonuclei (YF: 194 DEGs, OF: 1,234 DEGs), satellite cells (YF: 548 DEGs, OF: 2,784 DEGs) and FAPs (YF: 326 DEGs, OF: 2,102 DEGs). Satellite cells from OF demonstrated collagen signatures including elevated THBS1 expression and enrichment of TGF-β signaling. Concurrently, OF FAPs exhibited increased expression of fibroblast activation marker ADAMTS14. Furthermore, NicheNet analyses unmasked altered FAP-satellite cell communication in OF. Complementary *In vitro* experiments revealed that myogenic progenitor cells collected at 7d-recovery from OF (vs YF) donors displayed impaired myogenic differentiation and a cellular senescence-associated phenotype, while fibroblasts exhibited greater myofibroblast-like activation. Conditioned media derived from OF fibroblasts at 7d-recovery further increased cellular senescence and impaired myogenic potential (vs YF fibroblast conditioned media). Collectively, these findings suggest that recovery from disuse atrophy in older females is characterized by altered FAP-satellite cell communication and intrinsic function which may contribute to poor muscle remodeling.

**GRAPHICAL ABSTRACT:** Abstract Figure Legend
During re-ambulation following 14 days of unilateral limb immobilization, fibroblasts from older females (vs young females) exhibited an elevated activated myofibroblast-like phenotype. These changes were accompanied by altered fibro-adipogenic progenitor-satellite cell communication. Secretome derived from 7d-recovery fibroblasts increased cellular senescence and impaired myotube formation from older female donors. Collectively, these findings suggest that aging promotes a maladaptive fibrogenic secretory profile that impairs myogenic function and skeletal muscle remodeling following disuse atrophy.

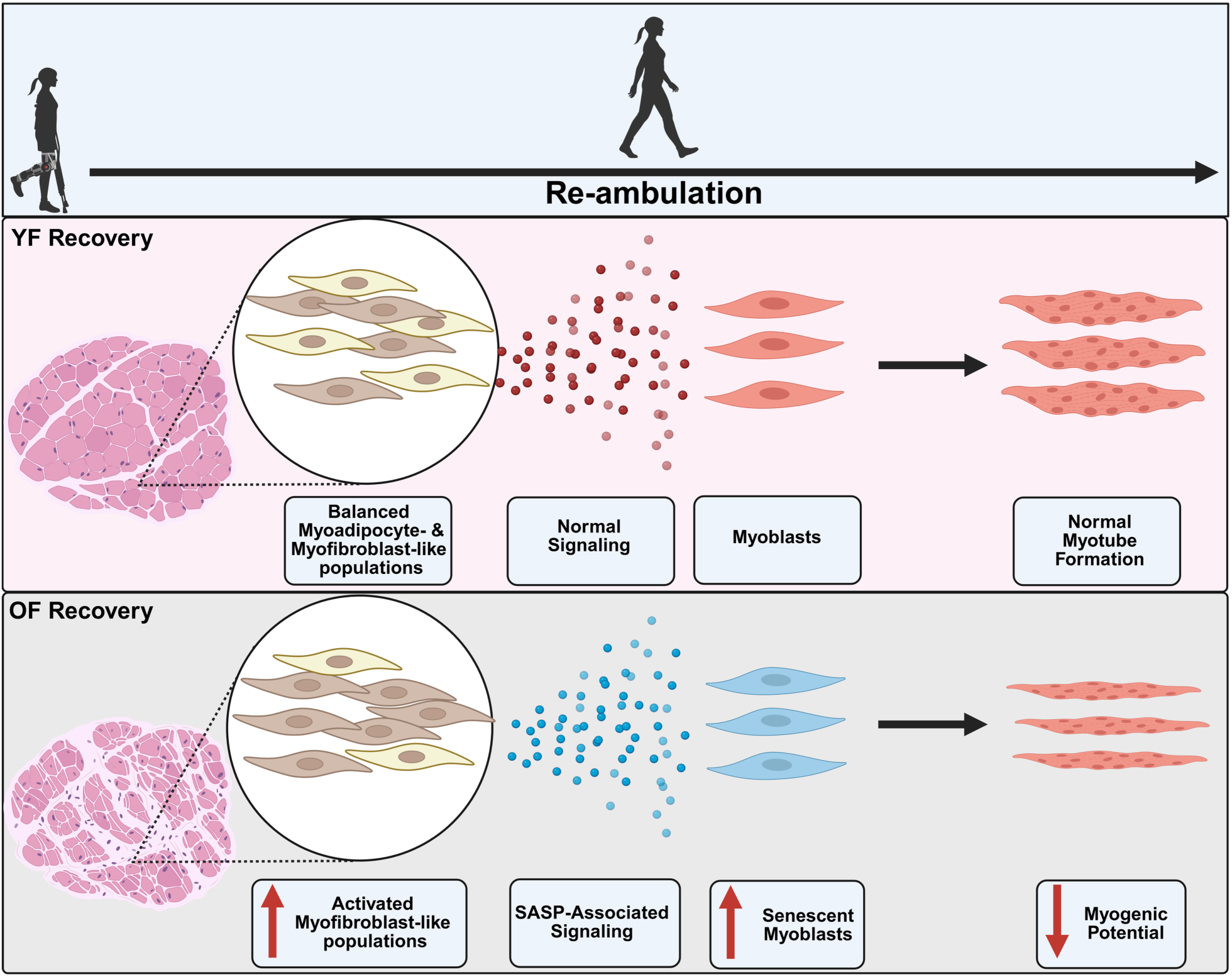

## INTRODUCTION

Skeletal muscle aging is characterized by muscle atrophy and functional decline ^1–3^. Contributing factors to impaired skeletal muscle health include disrupted function and intercellular communication among both myogenic (i.e., slow- and fast-twitch myofibers, satellite cells) and non-myogenic (endothelial cells, macrophages, fibro-adipogenic progenitors (FAPs)) cells ^4–9^. Additionally, older adults are particularly susceptible to periods of disuse-induced atrophy resulting from injury, illness, or hospitalization events and often exhibit poor muscle resilience, which increases risk of sarcopenia and consequently reduces mobility and quality of life ^10,11^.

Both myogenic and non-myogenic cell populations have critical roles in maintaining skeletal muscle health ^12–14^. For example, during muscle regeneration, satellite cells and FAPs are integral to the regenerative processes. Satellite cells activate, proliferate, and fuse to support muscle repair ^15^, whereas FAPs primarily differentiate into fibroblast- and adipocyte-like lineages and support muscle regeneration through interactions with satellite cells, extracellular matrix (ECM) remodeling, and coordination of tissue repair ^16^. Previous studies have shown that older adult skeletal muscle is characterized by reduced satellite cell pool and impaired myogenic function, which limits skeletal muscle regenerative capacity in response to muscle damage ^17–20^. Additionally, dysregulated FAP activation during aging can drive excessive collagen deposition, compromising skeletal muscle quality and contributing to muscle weakening ^21–23^. Emerging evidence further suggests that age-related accumulation of senescent cells may contribute to dysfunction in both myogenic and non-myogenic cell populations ^24–26^. Cellular senescence, characterized by cell-cycle arrest in response to cellular stressors such as DNA damage and oxidative stress, is elevated in aged skeletal muscle and may impair both myogenic and non-myogenic cellular functions and intercellular communication, and ultimately compromise muscle homeostasis ^27,28^. However, the molecular and cellular responses of satellite cells and FAPs, as well as the intercellular communication networks that underlie impaired muscle remodeling following periods of disuse in older adults, remain poorly understood.

Although both males and females face significant challenges in maintaining skeletal muscle health with aging, older women experience unique obstacles. While older men are more likely to develop sarcopenia, women with sarcopenia are more prone to severe disability and increased mortality risk ^29–31^. Additionally, Wu et al. reported that women display a greater rate of muscle atrophy than men during disuse ^32–34^. Older women also possess a smaller satellite cell pool than men, likely due in part to sex hormone differences ^35–37^, which may contribute to impaired regulation of satellite cell fate and inflammatory response following muscle damage ^38,39^. In a previous study, we observed dysregulated immune cell dynamics during the re-ambulation following 14d of limb immobilization, which presented an unique muscle remodeling response in the older females ^40^. Therefore, the purpose of this study was to gain further insight into the mechanisms of impaired muscle recovery in older females. To do so, we investigated the cellular responses during recovery following 14d of disuse-induced atrophy in young and older females using single-nucleus RNA-sequencing (snRNA-seq) and complementary cell culture approaches using participant muscle primary cells. We hypothesized that older women would be characterized by dysregulated myogenic and non-myogenic cellular responses, as well as intercellular signaling networks.

## METHODOLOGY

### Participant Demographics

Healthy, young (YF, n=8) and older females (OF, n=9) were included in this study based on findings from a previous analysis of the same cohort of young and older adults, which indicated that older females displayed an unique muscle remodeling response during recovery from disuse atrophy ^40^. As expected, groups differed in age (YF: 22±3 yrs, OF: 66±5 yrs; P<0.001) whereas no differences were observed in BMI (YF: 24±2 kg/m^2^, OF: 25±4kg/m^2^; P>0.05) or HbA1c (YF: 5.3±0.4%, OF: 5.5±0.3%; P>0.05). The study protocol was approved by the University of Utah Institutional Review Board (IRB #130232) and conducted in accordance with the *Declaration of Helsinki*.

Written informed consent was obtained from all participants prior to participation. The trial was registered at ClinicalTrials.gov (NCT04416191).

### Trial Design

Details of the trial design have been previously reported ^40^. Briefly, young and older females underwent 14 days of unilateral limb immobilization followed by 7d of re-ambulation recovery (**Fig 1**). During immobilization, participants wore a knee brace (Orthomen postop knee brace; Orthomen Inc., Foothill Ranch, CA, USA) on the dominant leg to prevent weight bearing and were provided crutches to assist with mobility. Skeletal muscle biopsies were obtained before and after immobilization and at 2d- and 7d-recovery for snRNA-seq and primary cell culture analyses.

**Figure 1.**
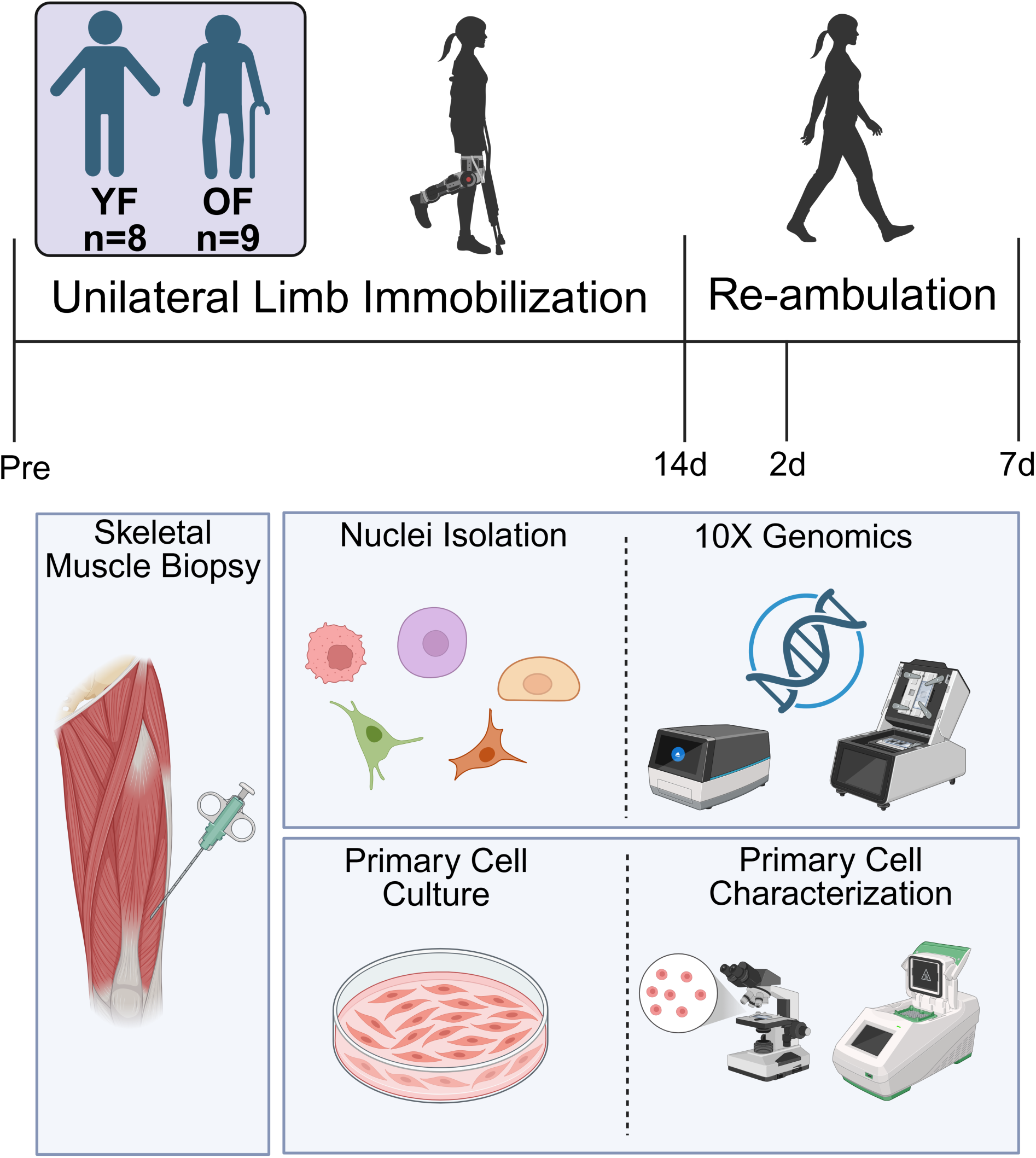
Trial schematic. Eight young and nine older females underwent 14d of limb immobilization followed by 7 days of re-ambulation. Vastus lateralis muscle biopsies were collected from the immobilized leg before (PRE) and after limb immobilization and on day 2 and 7 of re-ambulation. Muscle samples were processed for single-nucleus RNA sequencing or used for the isolation of myogenic progenitor cells and primary fibroblasts for *in vitro* cell characterization.

### Muscle Biopsies

As previously described ^40^, vastus lateralis muscle biopsies were obtained from the immobilized leg before (PRE) and after immobilization (POST), and at 2d- and 7d-recovery. Biopsies were collected using a 5mm Bergstrom needle with manual suction approximately 10-15 cm proximal to the patella. Subsequent biopsies were obtained ∼3 cm from the previous biopsy site. Muscle samples were either flash-frozen in LN2 and stored in −80°C for snRNA-seq or placed in Dulbecco’s Modified Eagle Medium (DMEM) supplemented with 1% penicillin and streptomycin (10,000 U/ml) for primary cell isolation.

### Single-Nuclei Isolation and RNA Sequencing

Three young (21±3 yrs) and older females (67±5 yrs) were identified for snRNA-seq analyses based on matching BMI values (YF: 25±3 kg/m^2^, OF: 23±4 kg/m^2^) and loss of total lean mass at post-immobilization (YF: −1.5±4.0%, OF: −2.2±1.4%).

Approximately, 20-30 mg of tissue from each biopsy was used for nuclei isolation. Single nuclei were isolated using the Chromium Nuclei Isolation Kit (10X Genomics, #1000494) according to the manufacturer’s recommendations. Briefly, frozen tissue was transferred to a pre-chilled sample dissociation tube and maintained on ice. Lysis buffer (200 µl) was added, and the tissue was minced with scissors and homogenized using a rubber pestle. An additional 300 µl of lysis buffer was then added, and samples were incubated for 10 minutes. The homogenate was transferred to a pre-chilled nuclei isolation column and centrifuged at 16,000xg for 20 sec at 4°C. Following filtration, nuclei were pelleted at 700xg for 3 minutes at 4°C. The pellet was resuspended in 500 µl Debris Removal Buffer and centrifuged at 700xg for 10 minutes at 4°C. The pellet was subsequently washed twice with Wash and Resuspension Buffer, with centrifugation at 700xg for 5 minutes at 4°C. Following the final wash, nuclei were resuspended in 60 µl of Resuspension Buffer, maintained on ice, and transported to the

University of Utah Genomics Core for sequencing. For each time point, the average total number of nuclei isolated collectively was 16,947 and 8,600 for YF and OF, respectively. snRNA sequencing was performed using the Genomics Chromium GEM-X Universal 3’ v4 Gene Expression 4-Plex Library and Expression Kit (10X Genomics). FASTQ files were aligned to the Human GRCH38-2024-A reference genome using Cell Ranger v9.0.0. to generate gene-barcode matrices ^41^. Ambient RNA was removed using CellBender v0.3.2 ^42^. Filtered matrices were processed in R with the Seurat v5.3.1 ^43^.

Nuclei with fewer than 750 features, more than 25% mitochondrial reads, or identified as doublets with scDBlFinder were excluded. Counts were normalized using SCTransform v2 ^44,45^. Datasets were integrated using the reciprocal PCA (RPCA) workflow in Seurat. Dimensionality reduction was performed using principal component analysis (PCA), and the first 30 principal components were used for UMAP visualization and clustering at a resolution of 0.5. Differential gene expression and cluster marker analyses were performed using the Wilcoxon Rank-Sum test. Functional enrichment analysis was conducted using Enrichr and Hallmark genes to identify overrepresented pathways and gene sets ^46^. Intercellular communication analyses were performed using CellChat and NicheNet according to the published workflow ^47^. The snRNA-seq dataset generated during this study has been deposited in the NCBI Gene Expression Omnibus (GEO) database under accession number GSE347901.

### Myoblast and Fibroblast Extraction

As previously described 30-50 mg of skeletal muscle tissue was minced and washed two times in Hank’s balanced salt solutions (HBSS without Ca^2+^, Mg^2+^) ^48^. Samples were then incubated for 30 minutes in a digestion cocktail containing collagenase II and trypsin. Digested muscle tissue was plated for 2h to allow fibroblast attachment. Media containing non-adherent cells were subsequently transferred to collagen-I-coated culture plate (Corning BioCoat Collagen-I). Cells were expanded in DMEM containing 5% FBS to passage 3 and cryopreserved in DMEM containing 5% FBS and 5% DMSO for subsequent phenotypic and functional assays.

### Cell Culture, Histology, and Immunofluorescence

For both myogenic progenitor cells (MPCs) and primary fibroblast experiments, four samples from four different participants within each group were pooled to generate YF (age: 23±3 yrs; BMI: 24±2 kg/m^2^) and OF (age: 65±5 yrs, BMI: 26±1 kg/m^2^) cell populations for each timepoint. For myogenic differentiation assays, MPCs were seeded in 6-well plates at 50,000 cells per well with growth media (GM; low glucose DMEM, 10% FBS). Once confluent, cells were then differentiated into myotubes using differentiation medium (DM; low glucose DMEM, 1% horse serum) for 6d. Myogenic function was subsequently assessed by quantifying the percentage of area occupied by myotubes and fusion index. For senescence analyses, MPCs were seeded in 12-well plates at a density of 40,000 cells per well with GM. Upon reaching approximately 50% confluency, cells were assessed for senescence-associated β-galactosidase (SA-β-gal) activity. Fibroblasts were plated in 6-well plates at 50,000 cells per well and assessed at approximately 70% confluency for lipid content (bodipy) and fibroblast activation (α-SMA^+^ TCF4^+^). Condition media were generated from fibroblasts collected at 7d-recovery in both the YF and OF groups. At the time of collection, media was centrifuged at 500xg for 5 minutes, and the resulting supernatant (conditioned media) was collected and stored at −80°C until further use. For conditioned media experiments, thawed conditioned media were applied to the PRE MPCs for 24 hrs before assessment of cellular senescence or initiation of myogenic differentiation.

Three to four randomly selected fields per well were Imaged by the Zeis Axioscan 7 fluoresence microscope (Zeiss, Jena, Germany) equipped with an X-Cite 120 LED Boost Illumination system (Excelitas Technologies, Waltham, MA, USA). Measurements from individual fields were averaged to obtain a single value for each well. Myogenic differentiation and lipid accumulation were quantified as the percentage of MF20^+^ area and bodipy^+^ area/cell, respectively, using FIJI following application of standardized intensity threshold. The percentage of SA-β-gal^+^ cells was calculated relative to the total number of cells using FIJI software. Fibroblast activation was assessed using Qupath by determining the percentage of α-SMA^+^ TCF4^+^ cells relative to the total number of cells.

Frozen OCT-embedded muscles samples were sectioned transversely at 10 µm using a cryostat (CM1860, Leica Biosystems, Wetzlar, DE). Muscle sections were stained to assess collagen turnover using B-CHP and collagen I labeling. Images were acquired using Zeis Axioscan 7 fluorescence microscope (Ziess, Jena, Germany) equipped with an X-Cite 120 LED Boost Illumination system (Excelitas Technologies, Waltham, MA, USA). Collagen turnover was quantified using FIJI software. Three randomly selected fields were analyzed per sample. Thresholds were applied to determine the percentage area positive for collagen-I and B-CHP staining within each field. Collagen turnover was calculated as the ratio of B-CHP^+^ area to Collagen I^+^ area (B-CHP/COL-I). Values from the three fields were averaged to obtain a single value for each sample. Complete histology protocols are provided in the Supplemental Materials and previous publications ^48–50^.

### RNA Isolation and RT-qPCR

Total RNA was isolated from PRE myotubes that were exposed to conditioned media derived from 7d-recovery fibroblasts using QIAzol lysis reagent (QIAGEN, #79306) (YF: n=6 replicates, OF: n=9 replicates). RNA extraction was performed according to the manufacturer’s instructions using chloroform phase separation followed by isopropanol precipitation. RNA pellets were resuspended in nuclease-free water, and RNA concentration was determined using an Epoch spectrophotometer equipped with a Take3 plate (BioTek, Winooski, VT). For cDNA synthesis, 0.4 µg of total RNA was reverse transcribed using the iScript cDNA synthesis kit (Bio-Rad, #17088-91) on a T100 Thermal Cycler (Bio-Rad) in a 20 µl reaction volume.

Quantitative real-time PCR (RT-qPCR) was performed using cDNA diluted1:2 in nuclease-free water on CFX Connect Real-Time PCR Detection System (Bio-Rad) with SsoAdvanced Universal SYBR Green Supermix (Bio-Rad, #317252-70). Gene expression was analyzed in duplicate and normalized to housekeeping gene GAPDH, and relative gene expression was calculated using the ΔΔCt (2^-ΔΔCt^) method. Target genes were selected to genes from the snRNA seq dataset and related to cellular senescence ^27^. Primers were purchased from Bio-Rad.

### Statistical Analysis

Statistical analyses were performed using GraphPad Prism, version 9 (GraphPad Software Inc., San Diego, CA, USA). Two-way ANOVA was used to determine main effects of age and time, with Sidak’s post-hoc test applied when significant interactions were detected. When missing values were present or groups were unbalanced, mixed effects models were used in place of ANOVA. T-tests were utilized to determine differences between groups in RT-qPCR gene expression. Data distributions were evaluated using Q-Q plots, and assessments of skewedness and kurtosis, with the Shapiro-Wilk testing performed when necessary. Statistical significance was accepted at P<0.05. Data are reported as mean±SD.

## RESULTS

### Cellular Profiling by snRNA-Sequencing

Using Seurat and Loupe Browser, nuclei were clustered according to their transcriptional profiles. Eight cell populations were identified including MHC I (slow-twitch) myofibers (MYH 7, TNNT1, TNNI1, MYL3, MYL2), MHC IIA (fast-twitch) myofibers (MYH2, TNNT3, TNNI2, ATP2A1), endothelial cells (VWF, PECAM1, CDH5), FAPs (PDGFRα, DCN, FBLN2), macrophages (CD68, CD163, MRC1), satellite cells (PAX7, MYF5), lymphocytes (T cells: CD3D, TRAC, LCK; natural killer cells: KLRB1, KLRD1, NKG7, GNLY), and mural cells (PDGFRβ, RGS5, ACTA2) (**Fig 2A-B**).

**Figure 2.**
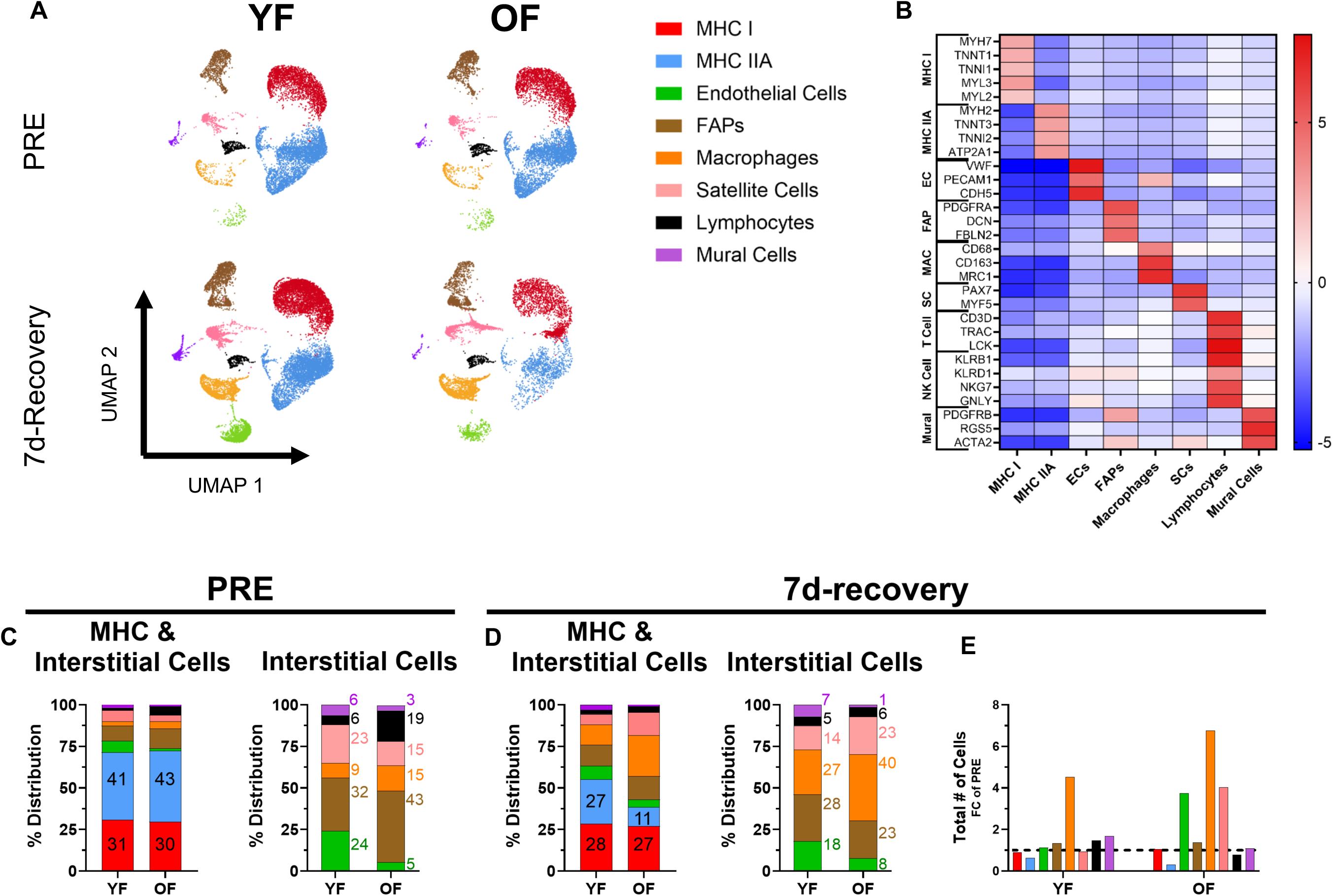
Single cell distribution in young and older females at baseline and at 7d-recovery. Nuclei isolated from skeletal muscle biopsies were clustered and annotated based on transcriptional profiles and visualized by UMAP (A,B). Cell-type distribution at baseline (Pre) (C) and 7d-recovery (D) and changes in cell abundance at 7d-recovery relative to PRE (E). EC, endothelial cell; FAPs, fibro-adipogenic progenitors; MAC, macrophage; MHC, myosin heavy chain; NK, natural killer; OF, older females; SC, satellite cell; YF, younger females.

Differences in cell population distributions between groups and changes in cells abundance relative to PRE are described in **Figure 2**. At PRE, the vast majority of nuclei were classified as slow-twitch (YF: 31%, OF: 30%) and fast-twitch (YF: 41%, OF: 43%) myofibers. Following removal of myofiber nuclei, FAPs represented the largest mononuclear population at PRE (YF: 32%, OF: 43%). YF and OF contained similar proportions of mural cells (YF: 6%, OF: 3%). In contrast, OF had lower proportions of endothelial cells (YF: 24%, OF: 5%) and satellite cells (YF: 23%, OF: 15%), alongside greater representation of immune populations, including macrophages (YF: 9%, OF: 15%) and lymphocytes (YF: 6%, OF: 19%) (**Fig 2C**). Minimal changes in cell-type proportions were observed at POST relative to PRE in either age group (**Fig S1A)**. At 2d-recovery, endothelial cells and lymphocytes were ∼2 fold more abundant in YF relative to PRE. In OF, most cell populations remained relatively unchanged at 2d-recovery relative to PRE, with the exception of lymphocytes which were ∼0.4 fold lower) (**Fig S1B**). By 7d-recovery, macrophages content appeared to increase in both groups relative to PRE (YF: ∼5 fold-change, OF: ∼7 fold-change). Additionally, both YF and OF contained lower numbers of fast-twitch nuclei (YF: ∼0.6 fold-change, OF: ∼0.3 fold-change). Endothelial cells and satellite cells were also present at ∼4 fold higher abundance in OF relative to PRE, whereas these populations appeared relatively unchanged in YF (**Fig2D-E**).

### Older Females Demonstrated a Robust Transcriptional Response Following Limb Immobilization

**Table 1** summarizes the number of differentially expressed genes (DEGs) identified within each cell population at POST and at 2-d and 7d-recovery relative to PRE. Relatively few DEGs were detected at POST (YF: 14, OF: 24). At 2d-recovery, YF exhibited substantially more DEGs than OF (YF: 813, OF 163), with most changes occurring in slow-(YF: 415, OF: 11) and fast-twitch (YF: 302, OF: 52) myonuclei. In contrast, OF demonstrated a markedly greater transcriptional response at 7d-recovery (YF: 1,548 DEGs, OF: 7,999 DEGs). The majority of these DEGs were identified in slow-twitch myonuclei (YF: 96, OF: 984), fast-twitch myonuclei (YF: 194, OF: 1,234), satellite cells (YF: 548, OF: 2,784), and FAPs (YF: 326, OF: 2,102).

**Table 1.**
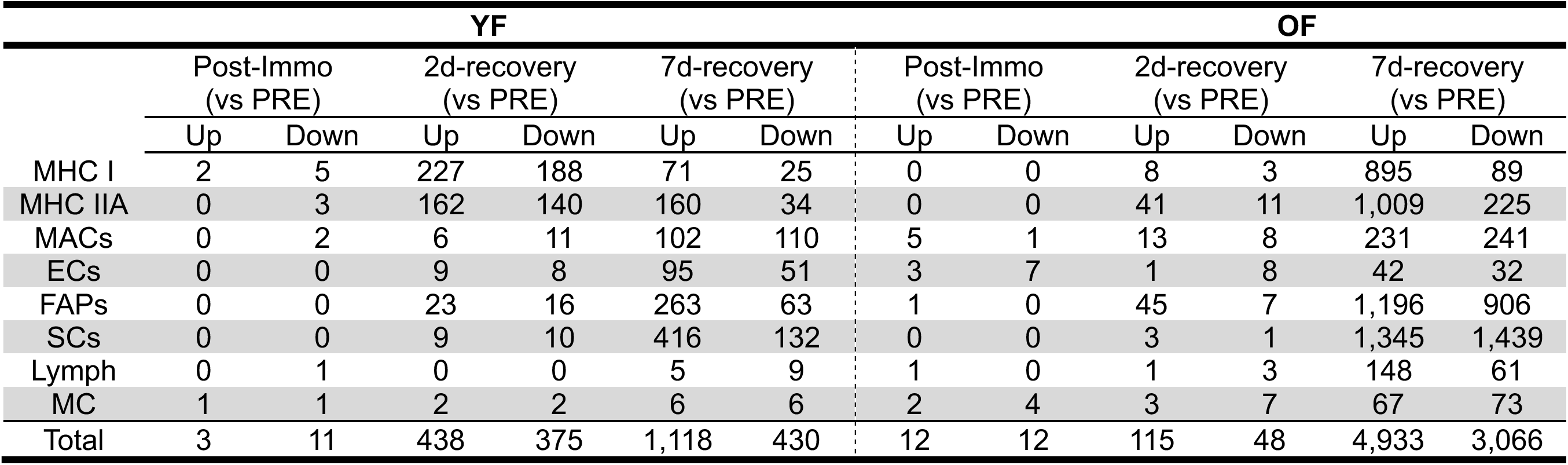
Number of DEGs at Post-Immobilization and during Recovery from Baseline.

|  | YF |  |  |  |  |  | OF |  |  |  |  |  |
| --- | --- | --- | --- | --- | --- | --- | --- | --- | --- | --- | --- | --- |
|  | Post-Immo<br>(vs PRE) |  | 2d-recovery<br>(vs PRE) |  | 7d-recovery<br>(vs PRE) |  | Post-Immo<br>(vs PRE) |  | 2d-recovery<br>(vs PRE) |  | 7d-recovery<br>(vs PRE) |  |
|  | Up | Down | Up | Down | Up | Down | Up | Down | Up | Down | Up | Down |
| MHC I | 2 | 5 | 227 | 188 | 71 | 25 | 0 | 0 | 8 | 3 | 895 | 89 |
| MHC IIA | 0 | 3 | 162 | 140 | 160 | 34 | 0 | 0 | 41 | 11 | 1,009 | 225 |
| MACs | 0 | 2 | 6 | 11 | 102 | 110 | 5 | 1 | 13 | 8 | 231 | 241 |
| ECs | 0 | 0 | 9 | 8 | 95 | 51 | 3 | 7 | 1 | 8 | 42 | 32 |
| FAPs | 0 | 0 | 23 | 16 | 263 | 63 | 1 | 0 | 45 | 7 | 1,196 | 906 |
| SCs | 0 | 0 | 9 | 10 | 416 | 132 | 0 | 0 | 3 | 1 | 1,345 | 1,439 |
| Lymph | 0 | 1 | 0 | 0 | 5 | 9 | 1 | 0 | 1 | 3 | 148 | 61 |
| MC | 1 | 1 | 2 | 2 | 6 | 6 | 2 | 4 | 3 | 7 | 67 | 73 |
| Total | 3 | 11 | 438 | 375 | 1,118 | 430 | 12 | 12 | 115 | 48 | 4,933 | 3,066 |

The number of DEGs comparing YF and OF at each timepoint are presented in **Table 2**. The total number of DEGs between groups increased from 224 at PRE to 1,837 at 7d-recovery. Most of these differences were observed in slow-twitch myonuclei (942), fast-twitch myonuclei (214), and satellite cells (651). Due to the robust transcriptional response at 7d-recovery, we decided to hone our attentions on this time point and specifically in myofiber, satellite cell, and FAP cell populations.

**Table 2.** Number of DEGs Identified in Older Females relative to Young Females.

|  | PRE |  | Post-Immo |  | 2d-recovery |  | 7d-recovery |  |
| --- | --- | --- | --- | --- | --- | --- | --- | --- |
|  | Up | Down | Up | Down | Up | Down | Up | Down |
| MHC I | 5 | 14 | 8 | 9 | 56 | 25 | 734 | 208 |
| MHC IIA | 116 | 57 | 171 | 71 | 60 | 110 | 69 | 145 |
| MACs | 0 | 1 | 3 | 5 | 3 | 1 | 0 | 1 |
| ECs | 11 | 1 | 3 | 1 | 26 | 10 | 8 | 18 |
| FAPs | 3 | 10 | 6 | 7 | 44 | 50 | 0 | 1 |
| SCs | 4 | 0 | 9 | 4 | 56 | 18 | 236 | 415 |
| Lymph | 0 | 2 | 0 | 0 | 1 | 0 | 0 | 0 |
| MC | 0 | 0 | 1 | 1 | 0 | 0 | 2 | 0 |
| Total | 139 | 85 | 201 | 101 | 246 | 214 | 1,049 | 788 |

### Slow- and Fast-twitch Myofiber Transcriptional Responses Following Limb Immobilization

We first examined shared and unique DEGs and enriched gene sets in slow- and fast-twitch myofibers at 7d-recovery. In slow-twitch myofibers, YF and OF shared 39 upregulated and 4 downregulated DEGs, whereas we observed 856 upregulated and 85 downregulated OF-specific genes (**Fig 3A**). There were also notable differences in the top 5 upregulated and downregulated DEGs between age groups. YF showed decreased expression of genes associated with myofiber structure and contractile function including NRAP and TNNT3, respectively. In contrast, OF demonstrated a decreased expression of HIF3A, suggesting reduced hypoxia-responsive signaling. Both age groups also showed evidence of altered metabolic regulation through distinct mechanisms. YF displayed reduced expression of PPP1R3C, a regulator of glycogen synthesis and storage, indicating potential suppression of glycogen metabolism, whereas OF showed decreased expression of GABP7, consistent with reduced fatty acid handling. Furthermore, both groups exhibited enrichment of genes involved in ECM and collagen remodeling, although the underlying transcriptional programs differed. YF preferentially upregulated structural ECM components involved in ECM organization and collagen assembly (HMCN1, MXRA5, FN1, GBN, and COL1A1), whereas OF preferentially upregulated genes linked to inflammation, collagen remodeling, extracellular signaling, and regenerative programs (POSTN, TNC, GPC6, S100A2, SFRP5) (**Fig 3B**). Despite these differences in DEG profiles, Gene Set Enrichment Analysis (GSEA) revealed similar pathway responses in both groups, characterized by enrichment of inflammatory response and epithelial-mesenchymal transition (EMT) pathways. However, OF also displayed upregulation of TNFα signaling via NF_K_B, another inflammatory pathway. Notably, YF slow-twitch myofibers demonstrated downregulation of several metabolic pathways, including oxidative phosphorylation, MyC Targets V1, MyC Targets V2, and adipogenesis, whereas no significantly downregulated pathways were detected in OF slow-twitch myofibers (**Fig 3C**).

**Figure 3.**
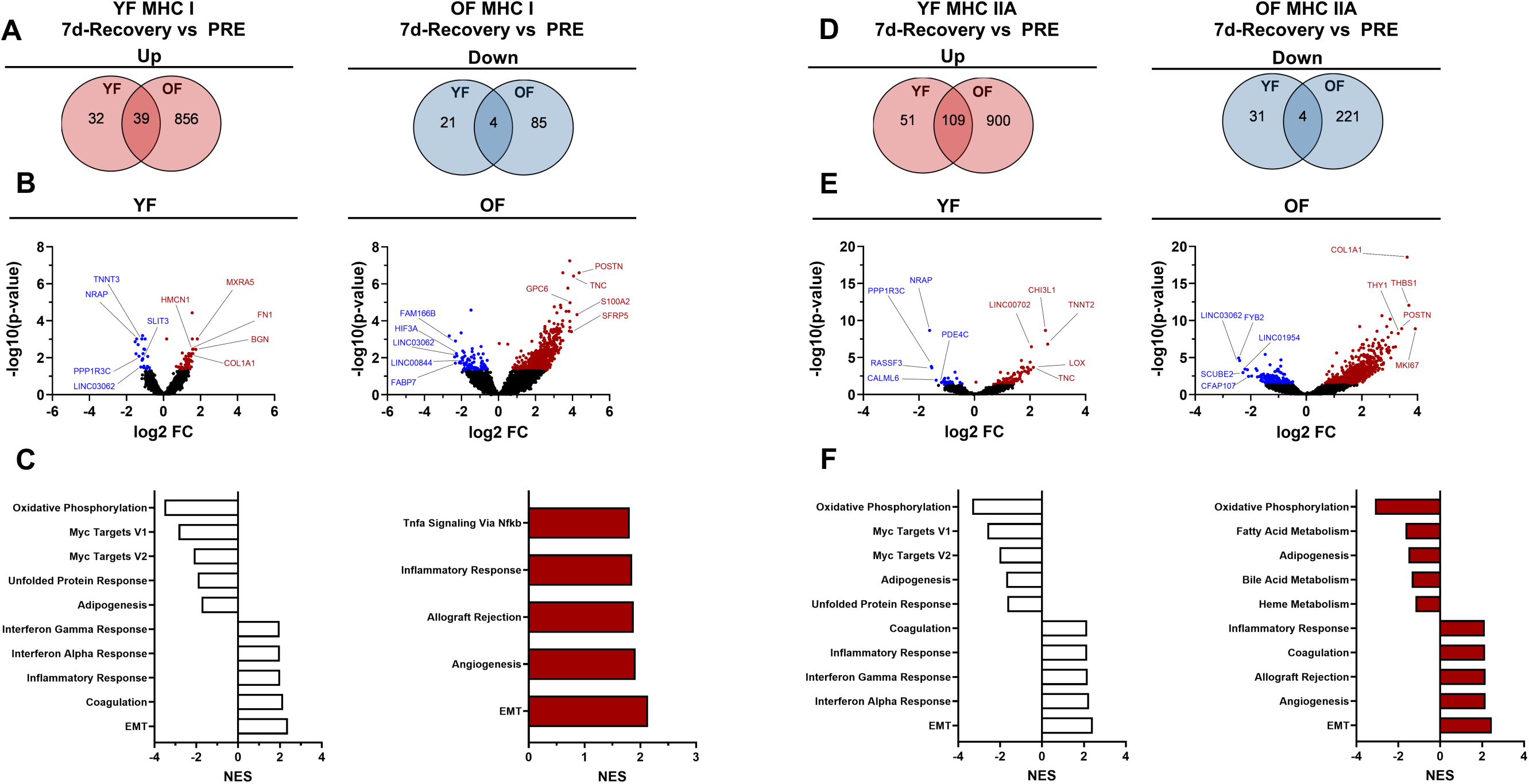
Age-related transcriptional response of slow- and fast-twitch myofibers during muscle recovery. Venn diagram illustrating the number of shared and unique upregulated and downregulated genes in slow-twitch myofibers at 7d-recovery relative to PRE in young and older females (A). Volcano plots of slow-twitch (MHC I) myofibers from young and older female at 7d-recovery relative to PRE, with the top 5 upregulated and downregulated genes labeled (B). Gene set enrichment analysis (GSEA) of slow-twitch myofibers showing the top 5 upregulated and downregulated pathways at 7d-recovery relative to PRE in young and older females (C). Venn diagram illustrating the number of shared and unique upregulated and downregulated genes in fast-twitch myofibers at 7d-recovery relative to PRE in young and older females (D). Volcano plots of fast-twitch (MHC IIA) myofibers from young and older females at 7d-recovery relative to PRE, with the top 5 upregulated and downregulated genes labeled (E). GSEA of fast- twitch myofibers showing the top 5 upregulated and downregulated pathways at 7d-recovery relative to PRE in young and older females (F). EMT, epithelial-mesenchymal transition; MHC, myosin heavy chain; OF, older females; YF, young females.

In fast-twitch myofibers, YF and OF shared 109 upregulated and 4 downregulated DEGs, whereas we observed 900 upregulated and 221 downregulated OF-related genes (**Fig 3D**). Additionally, fast-twitch myofibers exhibited age-dependent transcriptional remodeling. In YF, differential gene expression was characterized by reduced expression of genes associated with contractile structure and metabolic signaling (NRAP, PPP1R3C, PDE4C, CALML6) together with increased expression of genes involved in ECM organization and maturation (CHI3L1, LOX, TNC). In contrast, OF demonstrated a transcriptional profile indicative of collagen production and activated collagen remodeling characterized by increased expression of COL1A1, THBS1, POSTN, THY1, and MKI67 (**Fig 3E**). Similar to slow-twitch myofibers, fast-twitch myofiber pathway enrichment profiles were largely comparable between groups. Both YF and OF demonstrated downregulation of metabolic pathways (oxidative phosphorylation and adipogenesis), alongside upregulation of inflammatory pathways (inflammatory response and coagulation) and collagen-remodeling pathways (EMT) (**Fig 3F**). Together, these findings suggest that inflammatory and ECM remodeling pathways are common features of myofiber remodeling in young and older women following disuse atrophy. However, YF demonstrated greater collagen organization-associated signaling, whereas OF were characterized by a more pronounced tissue-remodeling transcriptional response within both slow- and fast-twitch myofibers.

### Older Females Demonstrate Impaired Satellite Cell Phenotype and Function During Muscle Recovery

In regard to satellite cell transcriptional response at 7d-recovery, YF and OF shared 95 upregulated and 321 downregulated DEGs, whereas we identified 1,344 upregulated and 1,024 downregulated OF-related genes (**Fig 4A**). Satellite cells from YF primarily exhibited a myogenic regenerative response characterized by increased RUNX1 expression and reduced expression of mature muscle structural genes (NRAP). In contrast, satellite cells from OF displayed reduced expression of genes associated with Wnt signaling (APCDD1) and ECM integrity (ITIH6), alongside increased expression of genes linked to ECM remodeling (THBS1, FRMD5) and stress responses (DMBT1, AHRR) (**Fig 4B**). GSEA revealed that both age groups demonstrated downregulation of metabolic pathways (oxidative phosphorylation and fatty acid metabolism). Both groups also displayed suppression of the myogenesis pathway and enrichment of EMT signaling at 7d-recovery. However, satellite cells from OF uniquely had enrichment of TGF-β signaling, suggesting enhanced collagen deposition pathways, along with upregulation of cell-cycle and proliferative-associated pathways (mitotic spindle, E2f targets, G2-M checkpoint). In contrast, YF displayed greater enrichment of inflammatory pathways (inflammatory response and interferon-γ response) and angiogenesis (**Fig 4C**). Due to the age-related differences in THBS1 expression, TGF-β signaling, and cell-cycle associated pathways, we sought to further characterize satellite cell phenotype and function using *in vitro* assessments.

**Figure 4.**
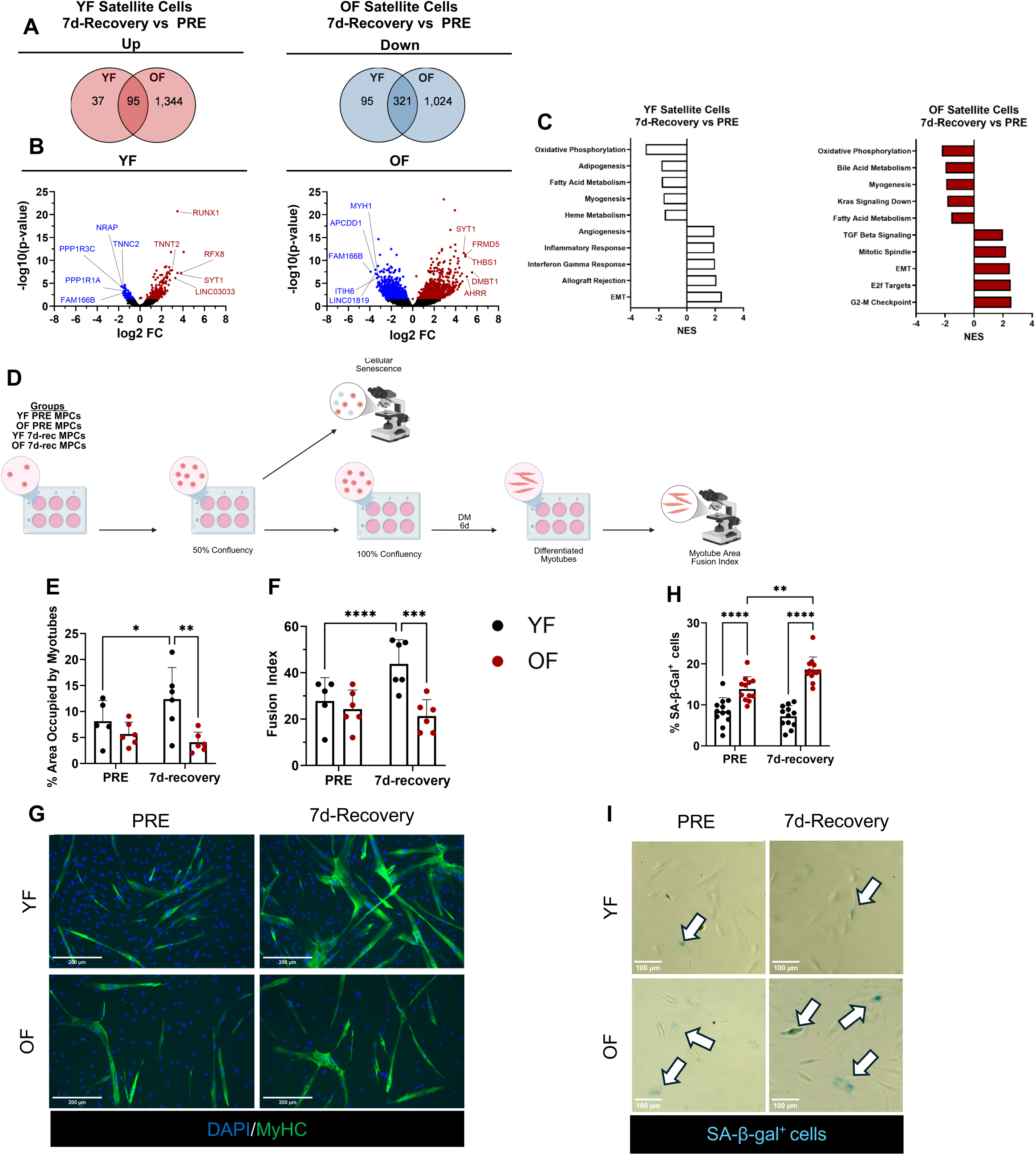
Impaired myogenic function in older females during muscle recovery. Venn diagram illustrating the number of shared and unique up and downregulate genes in satellite cells at 7d-recovery vs PRE in young and older females (A). Volcano plots of single nuclei transcripts from young and older female satellite cells at 7d-recovery relative to PRE, with the top 5 upregulated and downregulated genes labeled (B). GSEA of satellite cells showing the top 5 upregulated and downregulated pathways at 7d-recovery relative to PRE in young and older females (C). Schematic of the myogenic progenitor cell (MPCs) experimental design (D). Representative images and quantification of MPCs stained with MF20 and DAPI to assess % area occupied by myotubes and fusion index (n=6 replicates) (E-G). Representative images and quantification of MPCs stained with SA-β-galactosidase to assess cellular senescence (n=12 replicates) (H,I). Data are presented as mean±SD. DM, differentiation media; EMT, epithelial-mesenchymal transition; myogenic progenitor cells (MPCs) OF, older females; YF, young females.

Myogenic progenitor cells (MPCs) were differentiated and evaluated for myotube function (MF20) (**Fig 4D**). There were no group differences in myotube area or fusion index at PRE. However, YF exhibited a significant increase in myotube area and fusion index at 7d-recovery compared to PRE (myotube area: +6±5%, P=0.02; fusion index: +16±10%, P<0.01) and displayed greater myotube area (YF: 12±6%, OF: 4±2%; P<0.01) and fusion index (YF: 44±10%, OF: 21±7%; P<0.01) than OF at 7d-recovery (**Fig 4E-G**).

Due to the impaired *in vitro* myogenic potential observed in MPCs from OF, together with the increased expression of THBS1 and enrichment of TGF-β signaling in satellite cells from OF, both of which are associated with cellular senescence and stem cell aging ^51–53^, we hypothesized that satellite cells of OF may exhibit a senescence-associated phenotype. Consistent with this possibility and our previous observation that older adults from this cohort exhibited an elevated senescence response during recovery ^40^, we further evaluated satellite cell senescence. Notably, the snRNA-seq revealed that at the 7d-recovery, satellite cells from OF displayed a greater CDKN1A expression than satellite cells from YF (log2FC: 1.6, P<0.05; data not shown).

Therefore, we next assessed for senescence-associated β-galactosidase (SA-β-gal) positive cells in MPCs. At both PRE (YF: 8±3%, OF: 14±3%; P<0.01) and 7d-recovery (YF: 7±3%, OF: 19±3%; P<0.01), OF MPCs had a greater % of SA-β-gal^+^ cells than YF MPCs. Moreover, OF 7d-recovery MPCs demonstrated a greater senescent phenotype than OF PRE MPCs (+5±3%, P<0.01), whereas no change was observed in YF MPCs between the two timepoints (**Fig 4H-I**). Together, these findings suggest that satellite cells from OF adopt an active proliferative and collagen remodeling transcriptional program during recovery, accompanied by impaired myogenic function and cellular senescence *in vitro,* suggesting a potentially dysregulated regenerative response following disuse atrophy.

### Fibroblasts Exhibit Age-Related Phenotype Response to Muscle Recovery

FAPs from YF and OF shared 203 upregulated and 21 downregulated DEGs at the 7d-recovery relative to PRE, while OF exhibited 993 upregulated and 885 downregulated unique genes (**Fig 5A**). YF showed alterations primarily in cellular signaling and transcriptional regulatory pathways, characterized by increased expression of DGKI, ST6GAL2, and TMEM132D. In contrast, OF demonstrated a transcriptional profile consistent with collagen remodeling and mesenchymal differentiation, including increased expression of TGFBI, ADAMTS14, ALPL, and KANK4, together with reduced expression of the prostaglandin-catabolizing enzyme HPGD (**Fig 5B**). Both groups demonstrated downregulation of metabolic pathways (oxidative phosphorylation, fatty acid metabolism, and bile acid metabolism). EMT signaling was enriched in both groups, however, only YF had greater enrichment in TGF-β signaling and inflammation-related pathways (IL6 Jak Stat3 Signaling and Interferon-γ Response). In contrast, OF demonstrated stronger enrichment of cell-cycle and proliferation-related pathways (E2F targets, mitotic spindle, and G2-M checkpoint) (**Fig 5C**).

**Figure 5.**
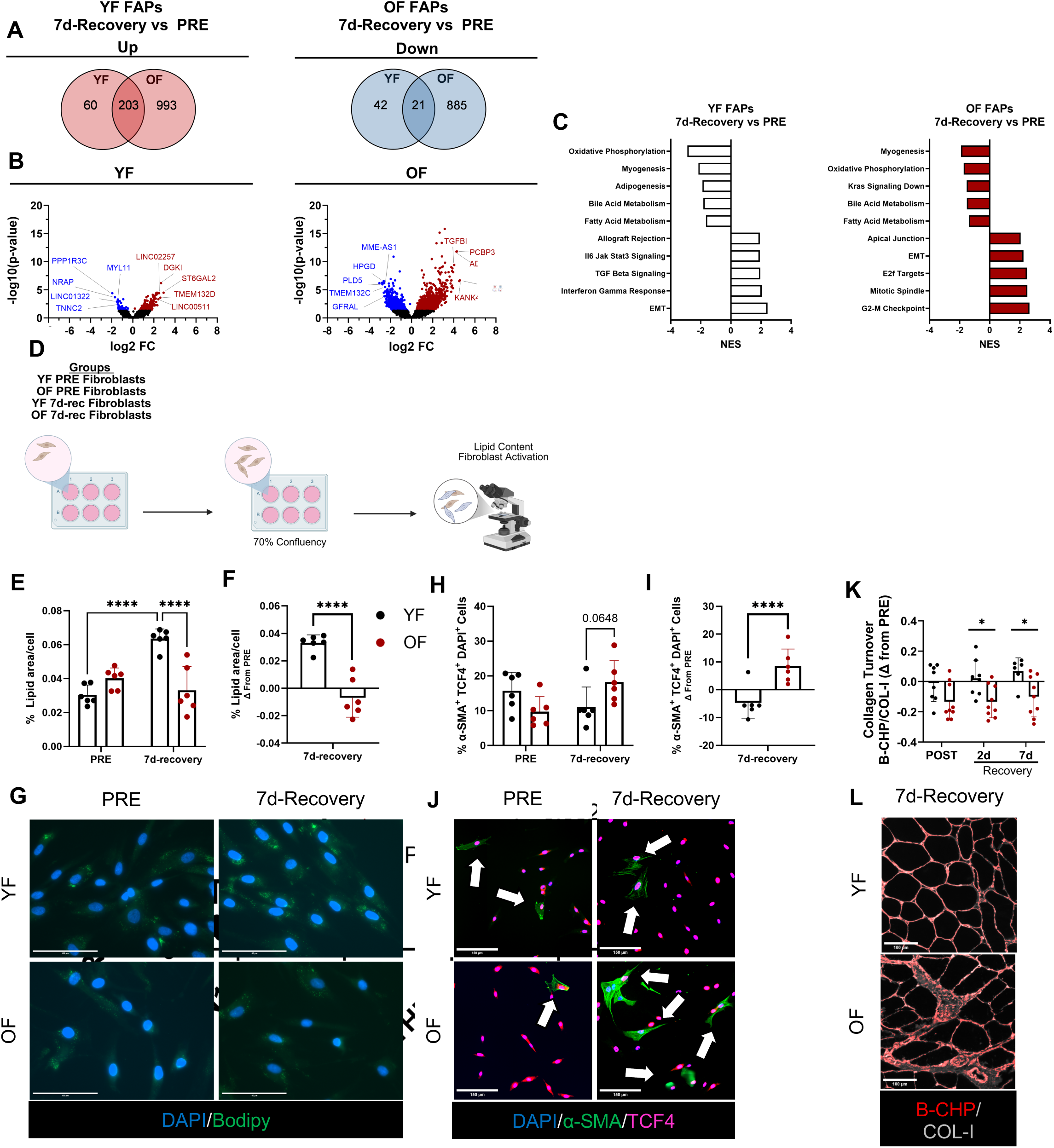
Older females exhibit greater fibroblast activation during muscle regrowth. Venn diagram illustrating the number of shared and unique upregulated and downregulated genes in FAPs at 7d-recovery relative to PRE in young and older females (A). Volcano plots of single nuclei transcripts from young and older female FAPs at 7d-recovery relative to PRE, with top 5 upregulated and downregulated genes labeled (B). GSEA of FAPs showing the top 5 upregulated and downregulated pathways at 7d-recovery relative to PRE in young and older females (C). Schematic of the primary fibroblast experimental design (D). Representative images and quantification of primary fibroblasts stained with bodipy to assess lipid content (n=6 replicates) (E-G). Representative images and quantification of primary fibroblasts stained with α-SMA and TCF4 to assess distribution of activated fibroblasts (n=6 replicates) (H-J). Representative images and quantification of collagen turnover in skeletal muscle cross-sections assessed by collagen hybridizing peptide (B-CHP) and collagen I (COL-I) (YF: n=8, OF: n=9) (K,L). Data are presented as mean±SD. EMT, epithelial-mesenchymal transition; OF, older females; YF, young females.

Given the upregulation of active collagen remodeling gene transcription observed in both slow- and fast-twitch myofibers, together with the fibroblast activation-associated expression of TGFBI and ADAMTS14 identified in OF FAPs, we sought to further characterize FAP phenotype *in vitro.* Primary fibroblasts were isolated to quantify lipid accumulation (bodipy) and myofibroblast activation (α-SMA^+^ TCF4^+^ cells) (**Fig 5D**). At PRE, lipid content did not differ between groups. However, YF had a greater increase in lipid area per cell at 7d-recovery relative to PRE compared to OF (YF: +0.033±0.006%, OF: −0.007±0.014%; P<0.01) and displayed greater lipid accumulation than OF at 7d-recovery (YF: 0.064±0.006%, OF: +0.033±0.014%; P<0.01) (**Fig E-G**). Similarly, no group differences were observed in the proportion of activated myofibroblasts (α-SMA^+^ TCF4^+^ cells) at PRE. However, OF displayed a greater increase in activated myofibroblasts at 7d-recovery relative to PRE than YF (YF: −4.7±5.8%, OF: +8.5±6.1%, P<0.01) (**Fig 5H-J**).

Because OF demonstrated a greater increase in activated myofibroblasts, we next assessed collagen remodeling in skeletal muscle samples within the larger cohort of YF and OF participants. Skeletal muscle cross-sections were stained for collagen I (COL-I) and biotin-conjugated collagen hybridizing peptide (B-CHP) to evaluate collagen abundance and collagen turnover. Compared with YF, OF exhibited greater collagen deposition (COL-I) following immobilization, with a significant increase at POST (YF: +2.9±2.3%, OF: +9.9±6.9%; P<0.01) and a trend toward greater collagen content at 7d-recovery (YF: +2.3±2.3%, OF: +7.2±5.6%; P=0.07) relative to PRE compared to YF (**Fig S2A-B)**. In contrast, YF displayed greater B-CHP staining at 2d-recovery from PRE compared to OF (YF: +1.5±2.0%, OF: −1.5±1.8%; P<0.01) (**Fig S2C-D)**, indicating increased collagen breakdown. Consistent with this finding, B-CHP to COL-I ratio (collagen turnover) was significantly greater in YF than OF at both 2- (YF: +0.021±0.119 B-CHP/COL-I, OF: −0.133±0.106 B-CHP/COL-I; P<0.01) and 7d-recovery (YF: +0.068±0.088 B-CHP/COL-I, OF: −0.100±0.134 B-CHP/COL-I; P<0.01) relative to PRE (**Fig 5K-L**). Collectively, these data indicate that fibroblasts from OF demonstrated a more activated myofibroblast-like population than those from YF, coinciding with reduced collagen turnover and elevated collagen accumulation during muscle recovery.

### Older Females Demonstrate Altered Intercellular Signaling During Muscle Recovery

Because coordinated cellular communication is critical for muscle regrowth, we examined intercellular networks using CellChat and NicheNet analyses. As a result, we found that the total number of inferred interactions was greater in OF at both PRE (YF: 273, OF: 331) and 7d-recovery (YF: 335, OF: 515). Given that satellite cells and FAPs demonstrated the largest transcriptional responses among mononuclear cell populations and that their intercellular communication is critical for effective muscle remodeling and often impaired in aged muscle ^23,54,55^, we focused on signaling interactions between these two cell types. When satellite cells were designated as receiver cells, NicheNet analysis predicted substantially greater FAP-derived regulation of targeted genes in OF than YF at 7d-recovery relative to PRE (**Fig 6A**). To determine whether fibroblast-derived factors influence MPC function and senescence-associated phenotype, conditioned media collected from fibroblasts at 7d-recovery were applied to PRE MPCs from the corresponding age group for 24hrs (**Fig 6B**). Relative to PRE MPCs maintained in standard growth conditions, treatment with fibroblast-conditioned media induced a trend towards reduced myotube size (YF: −0.5±2.6%, OF: −2.6±1.8%; P=0.09) and a significant impairment in myotube fusion index (YF: +1.4±4.5%, OF: − 16.0±3.8%; P<0.01) in OF relative to YF (**Fig C-E**). This response was accompanied by a greater increase in SA-β-gal^+^ cells in OF compared with YF (YF: +15±5%, OF: +27±7%; P<0.01) (**Fig 6F-G**). To further investigate senescence-associated pathways, RT-qPCR was performed on conditioned media-treated, MPC-differentiated myotubes for genes identified in the satellite cell snRNA seq dataset and genes associated with cellular senescence ^27^. MPCs treated with conditioned media from OF fibroblasts exhibited greater CDKN1A expression than those treated with conditioned media from YF fibroblasts (YF: 0.51±0.13, OF: 0.66±0.13; P<0.05). Additionally, expression of THBS1 (YF: 0.70±0.64, OF: 2.94±2.39; P=0.070), THBS2 (YF: 0.54±0.22, OF: 0.74±0.24; P=0.058), and TIMP2 (YF: 0.19±0.08, OF: 0.72±0.51; P=0.052) trended to be greater in OF compared to YF (**Fig 6H**). Together, these data suggest that FAP-satellite cell communication is altered during muscle recovery in OF and promotes senescence-associated phenotypes and impaired myogenic function.

**Figure 6.**
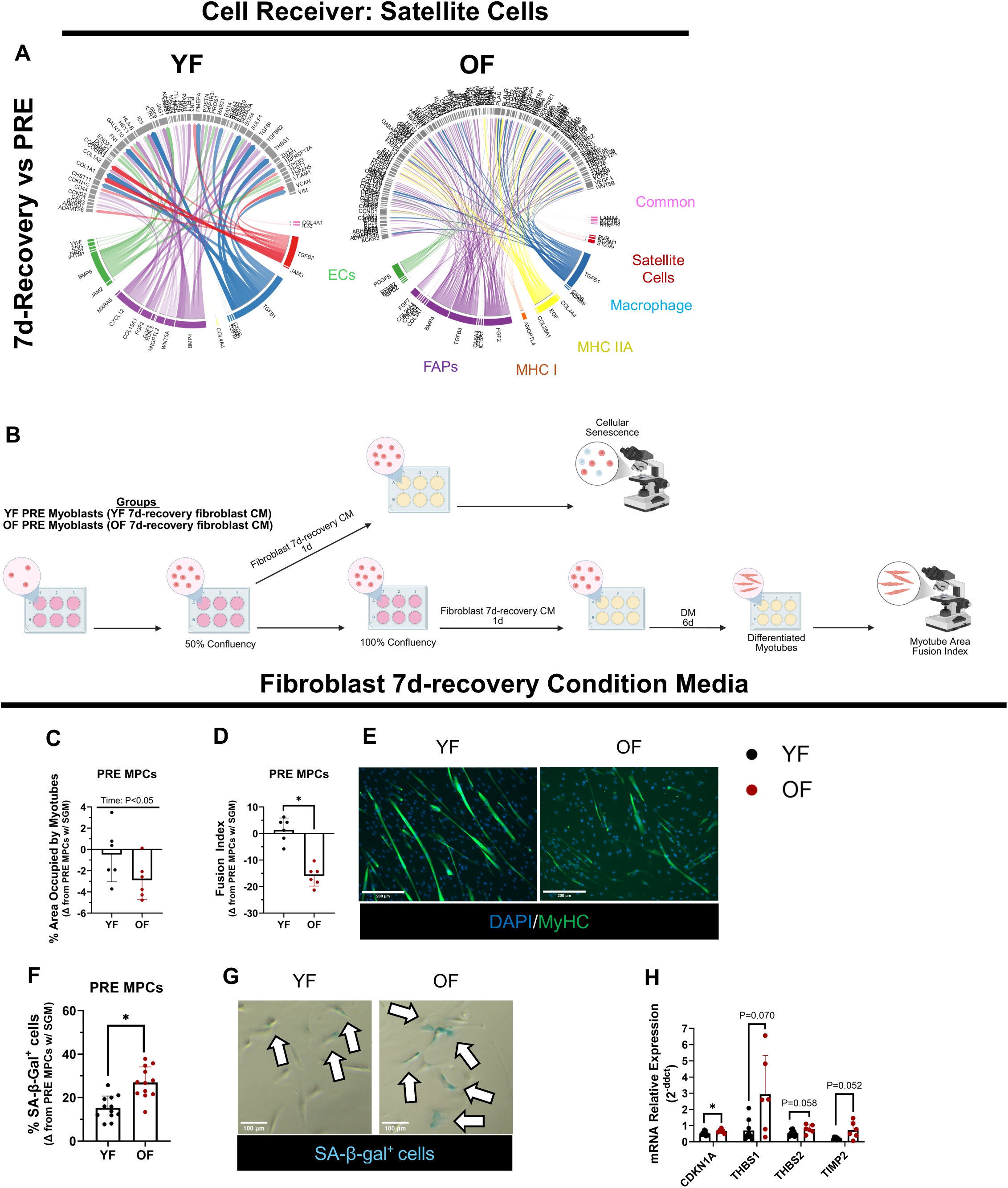
Altered fibroblast signaling disrupts myoblast function in older females during muscle regrowth. NicheNet analysis illustrating predicted intercellular signaling at 7d-recovery relative to PRE, with satellite cells designated as the receiver cell population (A). Schematic of the conditioned media experiment in which conditioned media from 7d-recovery fibroblasts were applied to PRE myogenic progenitor cells (MPCs) (B). Representation images and quantification of MPCs stained with MF20 and DAPI to assess % area occupied by myotubes and fusion index (n=6 replicates) (C-E). Representative images and quantification of MPCs stained with SA-β-galactosidase to assess cellular senescence (n=12 replicates) (F,G). RT-qPCR analysis of CDKN1A, THBS1, THBS2, and TIMP2 (YF: n=9 replicates, OF: n=6 replicates) (H). Data are presented as mean±SD. CM, conditioned media; DM, differentiation media; myogenic progenitor cells (MPCs); SGM, standard growth media; OF, older females; YF, young females.

## DISCUSSION

Here we investigated the skeletal muscle cellular and molecular responses to 14d of limb immobilization and subsequent recovery in young and older females. The major findings suggest that, during early re-ambulation from disuse atrophy, older females exhibit a robust transcriptional response in myonuclei, satellite cells, and FAPs that were indicative of a disrupted myogenic and fibrogenic phenotype and misaligned FAP-satellite cell communication. *In vitro* studies from primary cells of older female donors further confirmed increased myofibroblast activation, impaired collagen turnover, reduced myogenic function, and altered FAP-satellite cell communication, particularly at 7d-recovery, all of which may be related to dysregulated muscle remodeling in older females.

One of the primary findings of the present study was that older females demonstrated a substantially greater skeletal muscle transcriptional response during recovery, primarily driven by slow- and fast-twitch myonuclei, satellite cells and FAPs. Similar observations have been reported following acute eccentric muscle damage, where older adults displayed a greater muscle bulk transcriptional response than younger adults ^56,57^. These findings support that older adults have reduced transcriptional resilience in response to physiological stressors compared to younger adults. This markedly larger transcriptional response in older females may reflect unorganized and dysfunctional remodeling processes in effort to restore skeletal muscle homeostasis. Notably, satellite cells and FAPs from older females demonstrated enrichment of cell-cycle and proliferation-associated pathways, which may reflect increased cellular expansion and potentially contribute to the heightened regenerative response to modest muscle damage observed at 7d-recovery. This interpretation is consistent with our previous work demonstrating an increased number of central myonucleated fibers in older adults following immobilization ^40^. Together, these data support that re-ambulation following disuse atrophy elicits an exaggerated regenerative-like remodeling transcriptional response across myonuclei and several progenitor cells in older female skeletal muscle and akin to that observed following muscle injury and repair ^58–62^.

Interestingly, THBS1, one of the top upregulated genes in satellite cells from older females, has been identified as a key regulator of skeletal muscle mass and myoblast fate ^63,64^. Specifically, THBS1 expression is often elevated in aged muscle and activates TGF-β-SMAD2/3 signaling axis, which contributes to muscle atrophy through activation of the autophagy-lysosomal pathway and ubiquitin-proteasome system ^63^.

Additionally, elevated THBS1 expression has been shown to inhibit myoblast proliferation and differentiation ^64^, indicating that age-related expression of THBS1 may have detrimental effects on skeletal muscle remodeling in older females. THBS1 expression and TGF-β signaling have also been implicated in the induction of cellular senescence ^51,52,64^, which is consistent with our observation of greater SA-β-gal^+^ cells in MPCs during recovery from older females. At first glance, it appears contradictory that the myogenic cells from older females demonstrated an increased senescence-associated phenotype while simultaneously displaying enrichment of cell-cycle and proliferation-associated pathways. However, several mechanisms may explain this dichotomy. First, recent single-cell studies have demonstrated substantial satellite cell heterogeneity during regeneration, with proliferating, differentiating, and quiescence-returning satellite cell states coexisting within the same muscle regenerative environment ^65^. Therefore, the transcriptional profile observed in older females likely reflects a heterogenous population rather than a single, uniform satellite cell state which may have represented both senescent-associated and cell proliferative populations.

Furthermore, it is also possible that cellular senescence-associated pathways were upregulated to attenuate the regenerative response that promoted cell-cycle and proliferation pathways. Together, these findings suggest that satellite cells from older females activate proliferative programs during recovery while simultaneously exhibiting senescence-associated characteristics, indicative of dysregulated regenerative programming. Furthermore, MPCs from older females had impaired myotube formation during recovery. Previous studies have reported that aged muscle is characterized by increased cellular senescence and impaired myogenic function ^24,27,66,67^, and that cellular senescence can directly impair myotube formation and regeneration capacity ^25,68^. Taken together, these findings suggest that THBS1 and TGF-β signaling and cellular senescence may contribute to satellite cell dysfunction and impaired muscle recovery outcomes in older adults.

Another major finding from this study was the presence of an age-related FAP/fibroblast phenotype during recovery. Of note, FAPs from older females had greater transcriptional expression of TGFBI and ADAMTS14, genes associated with myofibroblast activation and profibrotic remodeling ^69,70^. This observation was consistent with our primary cell culture experiments showing that fibroblasts from older female donors at 7d-recovery displayed an intrinsic capacity to promote a greater proportion of activated myofibroblast-like cells, whereas fibroblasts from young females had greater adipogenic characteristics. These findings suggest that aging may shift fibroblast fate toward a more upregulated collagen remodeling phenotype during muscle regrowth. This interpretation is supportive of the finding in muscle cross sections indicating that older females had greater collagen accumulation and reduced collagen turnover during recovery relative to young females. During successful muscle regeneration, FAPs transiently adopt multiple differentiation states that support proper collagen remodeling, whereas persistent myofibroblast activation promotes dysregulated ECM remodeling and excessive collagen deposition ^71,72^. In other pathological conditions, including anterior cruciate ligament injury and cancer cachexia, expansion of FAPs and activated myofibroblasts have been implicated in muscle collagen deposition and fibrotic remodeling ^21,73^. Likewise, aged muscle is characterized by excessive collagen accumulation, particularly following muscle injury and periods of disuse atrophy ^27,40,74–78^. Collectively, these findings suggest that an age-related shift toward a more activated myofibroblast-like phenotype occurs alongside impaired ECM remodeling which may contribute to excessive collagen deposition that ultimately compromises muscle quality and muscle function ^79^.

Cellular communication and coordination are critical for successful muscle regeneration and regrowth following injury or disuse ^71,80–83^. Unfortunately, aging disrupts intercellular communication networks within skeletal muscle, which may contribute to impaired regenerative capacity and reduced muscle health ^27,84^.

Furthermore, the cellular secretome may directly influence the cellular crosstalk and muscle adaptation ^50,85^, highlighting the importance of paracrine signaling in regulating muscle regeneration. Consistent with these observations, another major finding of the present study was the altered FAP-satellite cell signaling during recovery in older females. This is in agreement with Thomas and colleagues who showed that coordinated communication among satellite cell, macrophages, and FAPs is a requisite for appropriate skeletal muscle adaptation and remodeling following mechanical overload but disrupted in aged mice ^86^. We also found that conditioned media from older female fibroblasts collected at recovery further promoted a cellular senescence fate in MPCs and impaired myogenic differentiation compared with conditioned media treatment from young female fibroblasts. These findings suggest that aging alters the fibroblast secretome during recovery which can have maladaptive downstream effects on MPC phenotype and function. Although the specific signaling factors responsible remain unknown, SASP molecules represent plausible candidates. Senescence cells secrete a variety of pro-inflammatory and pro-fibrotic mediators capable of inducing dysfunction in neighboring cells ^27,87,88^. While transient SASP signaling can facilitate tissue repair by recruiting immune cells and activating regenerative processes, persistent SASP production contributes to the chronic inflammatory environment associated with aging and impair muscle stem cell function ^89,90^. Therefore, it is possible that FAPs/fibroblasts cells from older females adopt a maladaptive secretory profile that disrupts pro-regenerative signaling, impairs myogenic potential, and contributes to poor muscle recovery.

Some limitations should be considered when interpreting these findings. First, the snRNA-seq analyses were performed in a subset of participants (n=3/group), which did not provide sufficient nuclei to identify subclusters within major cell types. Although snRNA-seq provides powerful cell-specific transcriptional information, its sensitivity remains limited and may not capture all age-related molecular responses that been observed using bulk transcriptomic approaches ^49^. Additionally, participants included in the snRNA-seq and primary cell culture analyses were not matched, and primary cells were pooled for functional experiments, limiting our ability to directly relate transcriptional signatures to functional cellular and morphological muscle outcomes.

Nevertheless, the integration of snRNA-seq, primary cell culture, and histological analyses provide a unique framework for investigating cellular mechanisms underlying impaired muscle recovery with aging.

In summary, we surmise that during muscle recovery from disuse atrophy, FAPs/fibroblasts from older females exhibit an age-related transcriptional and functional responses characterized by increased myofibroblast activation and altered intercellular signaling that impairs myogenic function and cellular remodeling. Collectively, these findings identify FAPs/fibroblast-driven signaling networks as potential therapeutic targets for improving muscle remodeling from disuse in older adults.

## Author Contribution

Conceived and designed research: MD and RO

Experiments and data analysis: CS, AK, MD, CF

Analyzed and interpreted results of experiments: CS, ZF, PB, EY, RC, AK, CF, MD

Publication preparation: CS, MD

Manuscript editing: all authors

## Acknowledgements

We thank the participants for their dedication and commitment to this research. We also thank the Clinical and Translational Science Institute nursing and medical staff for their assistance with muscle biopsies, blood draws, and participant care. Additionally, we thank Dr. Allen Opal (University of Utah Genomics Core) for her assistance with single-nuclei RNA sequencing and Dr. Chris Stubben (University of Utah Bioinformatics Core) for his assistance with the bioinformatic analyses of the single-nuclei RNA sequencing dataset.

## Supplemental Information

Supplemental Methodology

Supplemental Figures S1-2

## Supplemental Methods

### Immunocytochemistry and Histology

#### SA-β-galactosidase

SA-β-galactosidase Cell Technology Kit (#9860) was used to detect senescence in the primary myoblasts and fibroblasts. Cells were fixed with fixative solution for 15 minutes. After a few rinses with 1xPBS cells were incubated with β-galactosidase staining solution (pH=6.0) overnight. On the next day the β-galactosidase was removed and rinsed with 1xPBS and then preserved for long-term storage in 70% glycerol and stored at 4°C.

#### MF20

Cells were rinsed 3× with 1xPBS and then fixed with 4% PFA for 15 minutes. Cells were then permeabilized with 0.1% Triton X-100 in 1xPBS for 15 minutes. were washed (3×5 minutes) in 1xPBS and then blocked with 2% BSA for 60 minutes at RT. After 1 rinse with 1xPBS cells were incubated with 1° MF20 (MF20-DHSB, 1:100 in 2% BSA) overnight at 4°C. On the next day cells were washed (3×5 minutes) with 1xPBS and incubated with 2° antibodies (AlexaFlour 488 H/L, 1:250) for 60 minutes at RT. Then after the cells were washed (3×5 minutes) in 1xPBS cells were incubated with DAPI (1:10,000) for 10 minutes. Lastly, cells were rinsed 3× with 1xPBS and leavingPBS in the wells for imaging.

#### Bodipy

Cells were washed (3× 3 minutes) in 1xPBS and then fixed with 4% PFA for 20 minutes at RT. After washes (3×3 minutes) 1xPBS cells were incubated with Bodipy (10 ug/ml, ThermoFisher, #D3922) for 1hr. Then cells were washed (3×3 minutes) in 1xPBS and incubated with DAPU (1:10,000) for 10 minutes. Lastly, cells were washed (3×3 minutes) in 1xPBS and leaving PBS in the wells for imaging.

#### α-SMA and TCF4

Cells were rinsed with 1xPBS and fixed with 4% PFA for 20 minutes. After washes (3×3 minutes) in 1xPBS cells were permeabilized with 0.1% Triton-X100 in 1xPBS. Then cells were blocked with 1% BSA + 0.1% Triton for 1hr. Then cells were incubated in 1° TCF4 (Cell Signaling, #2569, 1:100) and α-SMA (Santa Cruz, #sc-130616, 1:100) in 1% BSA + 0.1% Triton for 2hrs at RT. After washes (3×3 minutes) in 1xPBS + 0.1% Triton. Cells were incubated in Gt α Rb Biotinylated 2° antibody (Jackson Immuno Research, #111-065-003, 1:1000) and Gt α Ms IgG2a AF488 (Life Technologies, #A-21131, 1:250) in 1% BSA + 0.1% Triton for 85 minutes. After washes (3×3 minutes) in 1xPBS + 0.1% Triton cells were incubated in SA-HRP (1:100) in 1xPBS + 0.1% Triton for 1 hr. Cells were then washed (3×3 minutes) in 1xPBS + 0.1% Triton and incubated with AlexaFluor 555 amplification (ThermoFisher, #B40933, 1:200) in 1xPBS + 0.1% Triton for 20 minutes. Cells were washed (3×3 minutes) in 1xPBS + 0.1% Triton and incubated with DAPI (1:10,000) in 1xPBS + 0.1% Triton. Lastly, cells were washed (3×3 minutes) and prepared for imaging.

#### COL-I/B-CHP

Muscle sections were air dried for 1hr at RT and then fixed with chilled acetone for 20 minutes. After 3×3 minutes 1xPBS washes, sections were blocked with 2.5% normal horse serum for 1hr at RT. Afterward sections were blocked with streptavidin (4 drops/ml of 2.5% normal horse serum) and biotin (4 drops/ml of 2.5% normal horse serum) for 15 minutes each. In preparation for incubating the sections with B-CHP (15 µM in 2.5% normal horse serum) the B-CHP solutions was warmed with aheating block (80°C) for 5 minutes and then quickly cooled in ice water for 2 minutes and then collagen-I (Abcam, #34710, 1:100) was added to the B-CHP (15 µM) and quickly added to the section and left overnight at 4°C. On the next day the sections were washed 3×5 minutes in 1xPBS and incubated with Dylight (ThermoFisher, #SA-5549, 1:200) and 2° AlexFlour 647 (Gt α Rb IgG, Invitrogen, #21245, 1:250) for 1 hr at RT. Lastly slides were washed 3×3 minutes in 1xPBS and coverslips were mounted with VectorShield mounting media.

---

**Figure S1.**
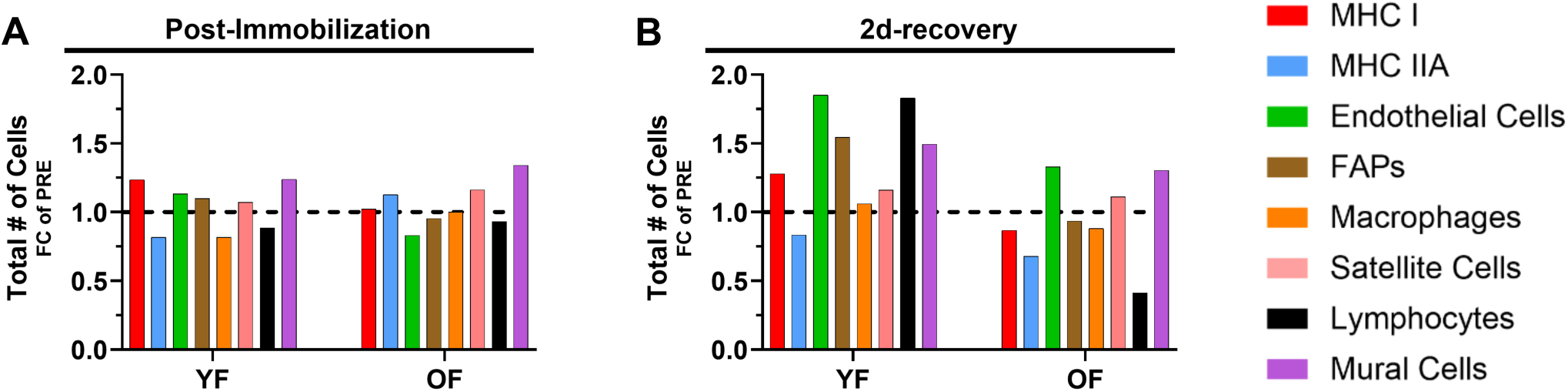
Change in total number of cells at post-immobilization and 2d-recovery. Changes in cell abundance at post-immobilization and 2d-recovery relative to PRE (A,B). FAPs, fibro-adipogenic progenitors; MHC, myosin heavy chain; OF, older females; YF, younger females.

**Figure S2.**
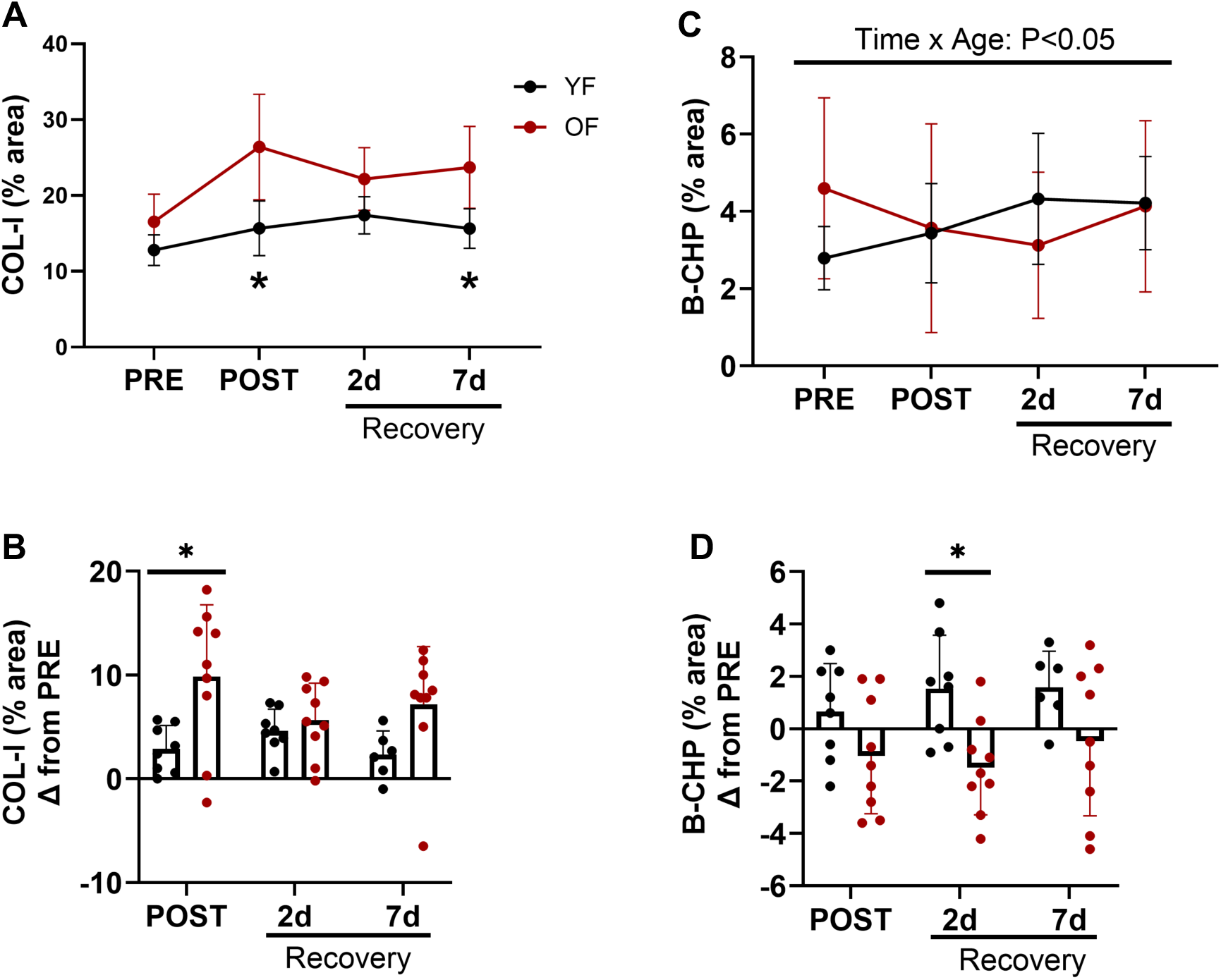
Collagen remodeling after immobilization and during recovery. Quantification of collagen deposition (% COL-I) and collagen hybridizing peptide (% B-CHP) in skeletal muscle cross-sections (A-D).

